# Climate change shifts abiotic niche of temperate skates towards deeper zones in the Southwestern Atlantic Ocean

**DOI:** 10.1101/2021.02.17.431632

**Authors:** Jéssica Fernanda Ramos Coelho, Sergio Maia Queiroz Lima, Flávia de Figueiredo Petean

## Abstract

Climatic changes are disrupting distribution patterns of populations through shifts in species abiotic niches and habitat loss. The abiotic niche of marine benthic taxa such as skates, however, may be more climatically stable compared to upper layers of the water column, in which aquatic organisms are more exposed to immediate impacts of warming. Here, we estimate climate change impacts in Riorajini, a tribe of four skates, as a proxy to (1) evaluate the vulnerability of a temperate coastal zone in the Atlantic Southwest, and (2) study niche dynamics in a scenario of environmental changes on this group of threatened species. We modelled each species abiotic niche under present (2000–2014) and future (2100, Representative Concentration Pathway 8.5) climatic scenarios, then measured niche overlap, stability, expansion, and unfilling. Our results reveal an expansion of suitable environment for the occurrence of the tribe in up to 20% towards deeper areas (longitudinal shift), although still within the limits of the continental shelf. We discussed the downfalls of such shift to the species and to the local biota in newly invaded areas, and suggest that even deeper layers of marine temperate zones are vulnerable to dramatic environmental changes as a consequence of global warming.

## INTRODUCTION

Long-term analyses show that global warming resulting from the continuous increase in emission of greenhouse gases is likely an effect of anthropogenic activities (Houghton 1996; Mann et al. 1999; Barnett et al. 2001; Rosenzweig et al. 2008). Such changes are occurring faster than most organisms can adapt to (Quintero and Wiens 2013), and, besides the difficulty to attribute an impact as a consequence of anthropic global warming, studies are consistently finding rather compelling evidence of theoretical predictions for climate change-related impacts on biodiversity distribution (Hughes 2000; Walther et al. 2002; Parmesan and Yohe 2003; Perry et al. 2005; Chivers et al. 2017). Terrestrial organisms are changing their distributions to higher latitudes and elevation to cope with thermal stress (Root et al. 2003; Hickling et al. 2006; Chen et al. 2011), while in the oceans these differences are latitudinal but also in depth (Perry et al. 2005; Nicolas et al. 2011). Fewer physical barriers in comparison to terrestrial habitats make it easier to move and disperse in the marine environment (Pinsky et al. 2013), although some biological and ecological characteristics can constrain populations’ ability to escape/avoid these adverse events (Somero 2010).

Studies on the distribution of species under different geographic, temporal and climatic scenarios benefited from the increasing availability of biodiversity data in online databases (e.g., GBIF; https://www.gbif.org) coupled with recent methodological frameworks and computational models able to address biological issues (e.g., Guisan and Zimmermann 2000; Broennimann et al. 2012; Guisan et al. 2014). These advances boosted our capacity to test ecologic hypothesis and visualize theoretical scenarios more efficiently, with a growing body of literature using models to estimate regions of potential occurrence of species, to indicate ecological barriers to populations, and identify vulnerable areas to climate change impacts (Watson et al. 2013; Costa et al. 2017). However, perhaps because of sampling difficulties and costly logistics, studies focusing on marine taxa are yet scarce in comparison to terrestrial ones (Dambach and Rödder 2011), and from those published so far applying ecological niche models (ENMs) for marine taxa, over half focus on groups like bony fish, mollusks, and marine mammals (Melo-Merino et al. 2020), while elasmobranchs are, to some degree, neglected. For threatened species, the use of such non-invasive methodologies is of paramount importance to provide the basis from which conservation efforts can be planned and implemented (Hammerschlag & Sulikowski 2011).

Due to global warming, large-scale changes in environmental conditions shift species abiotic niches besides reshaping communities with unprecedented impacts on their biodiversity. A species’ abiotic niche, also called Grinnellian niche, comprises the set of abiotic and climatic optimal conditions in which populations occur (Soberón & Nakamura 2009). In marine ecosystems, the most evident impacts of global warming on species’ abiotic niche include an increase in mean temperature, rise in sea level, ocean acidification, and a decrease in levels of dissolved oxygen (Nicholls et al. 2007). The “tropicalization” of coastal marine temperate communities, for example, illustrates a shift of biodiversity distribution consequent of these changes, with an expansion of warm-water species dominance that has direct economic implications (e.g., changes in fisheries global catch; Cheung et al. 2013) and tremendous ecological impacts (e.g., food-web interactions; Harley 2011). The accumulation and synergistic effect of climate-related changes can alter abiotic conditions of the planet to the extent that living organisms will be pressured towards the maxim “move, adapt or die”.

For elasmobranchs, animals that generally have long life cycles, slow growth, and late maturation, the “move” option seems more feasible in face of climatic stresses (Stevens et al. 2000; Helfman et al. 2009). Skates, however, elasmobranchs that lay sessile egg-capsules, tend to show philopatry with strong reliance on particular shallow-water habitats to reproduce, which increases the vulnerability of this group to climate changes in comparison to other pelagic elasmobranchs (Dulvy and Reynolds 2002; Parmesan 2006; Dulvy et al. 2014; Di Santo 2015). Besides, a geographic expansion beyond the limits of the continental shelf would require those strictly coastal skates to evolve complex adaptations, such as in osmoregulatory functions (Treberg and Speers-Roesch 2016), unfeasible in a short period of time in which climate is changing. On the other hand, because skates are mainly benthic species, occurring in deeper layers of the water column, their abiotic niche may be more stable compared to organisms that occur in superficial layers, where the effects of an increasing heat are immediate (e.g., Pearce & Feng 2013). Therefore, considering ecological characteristics (benthic, sedentary habit) and putative climate change impacts on coastal zones, tracking skates’ response to climate change is a good indicator of the extent of environmental change and climatic instability at coastal zones.

Here, we model the response of Riorajini’ skates to climate change as a case study to estimate its impacts in a temperate coastal zone. Riorajini (*sensu* McEachran and Dunn 1998) is a clade of four Arhynchobatidae skates occurring in sympatry in the Southwest Atlantic Ocean, a region that harbors the highest number of threatened chondrichthyan species in the Neotropics (Field et al. 2009). According to the IUCN latest global assessment, *Atlantoraja castelnaui* (Miranda Ribeiro, 1907) is evaluated as ‘endangered’, while *A. cyclophora* (Regan, 1903), *A. platana* (Günther, 1880), and *Rioraja agassizii* (Müller & Henle, 1841) are ‘vulnerable’ (Hozbor et al. 2004; Massa et al. 2006; Kyne et al. 2007; San Martín et al. 2007). These species occur mainly at the Warm Temperate Province (Spalding et al. 2007), influenced at north by the Cabo Frio upwelling system, and at south by the effects of cold-water masses from the Malvinas current (Peterson and Stramma 1990; Coelho-Souza et al. 2012). Since Riorajini species exhibit signs of conserved abiotic niches (Coelho et al. 2020), they may track changing environmental conditions more closely given their tendency to maintain ancestral lineages’ characteristics of niche (Chivers et al. 2017). Thus, these species may shift their geographic distribution if climatic changes interfere in abiotic dynamics at the temperate zones where they occur, given probable loss of cold-water habitats.

We hypothesize Riorajini species will present a southward shift in their current geographic distribution to avoid thermal stress in a future scenario of global warming, with potential reduction in areas where they are likely to occur as the environment gets warmer. To test this hypothesis, we used compiled information from public biodiversity databases to compare dynamics of abiotic niches between models of current (2000– 2014) and future (year 2100) distributions of the four Riorajini species. If held valid, our hypothesis suggests that benthic layers of coastal zones are vulnerable to global warming impacts similarly to upper oceanic layers.

## MATERIAL AND METHODS

We coupled ecological niche models (ENMs) with an ordination approach to visualize and measure the degree of change between niches of present and future climatic scenarios for each species, following the methodological framework of Broennimann et al. (2012). We ran the models using MaxEnt in the dismo package (Hijmans et al. 2017) in R 3.5.1 (R Core Team 2018). We chose this machine-learning algorithm because it outperforms others when measuring niche overlap between native and invaded areas (Broennimann et al. 2012), which is one of our goals. Maps were edited using QGIS 2.8.9.

### Models of present and future climatic scenarios

To visualize and characterize (in terms of variable importance) the niche of each Riorajini skate species in two different climatic scenarios, ENMs followed a correlative approach between georeferenced sites of occurrence of each taxon, and data characterizing climatic conditions where such is present (also called ‘abiotic predictors’, or ‘layers’) (Phillips et al. 2006; Robinson et al. 2011). We used the dataset of Coelho et al. (2020), which compiled occurrence records for the four Riorajini species from published literature and public online databases. These records were filtered to include only specimens preserved in ichthyological collections or museums, collected in the past 75 years, because older records are often inaccurate (Zizka et al. 2020), and georeferenced within the area of known occurrence of the group to increase data reliability.

To model future climatic scenarios, we used data from Representative Concentration Pathways (RCPs). RCPs represent data from the literature on possible paths for the main driving agents of climate change. There are currently four RCPs available, from mild to more extreme scenarios, varying from 2.6 to 8.5 W/m^2^ (ranging from ∼490 to ∼1370 ppm CO_2_, respectively) predicted from the trends in emission of greenhouse gases and land use for the end of the century (Van Vuuren et al. 2011). These emissions translate into an increase of up to 1.7 °C in mean temperature in the 2.6 RCP scenario, and up to 4.8°C in the 8.5 RCP scenario, both compared to preindustrial levels (Stocker et al. 2013). We used layers for future (predictions for year 2100) environmental conditions considering the worst-climatic-scenario (RCP 8.5) for which 18 variations (e.g., minimum, maximum, mean) of three variables (salinity, temperature and currents velocity) for benthic maximum depth were available in Bio-ORACLE (Tyberghein et al. 2012; Assis et al. 2017). We considered the RCP 8.5 as the most probable reflection of future impacts of climate change given current business-as-usual practices; thus, such catastrophic scenario may also be a cautious approach to estimate the impacts of ongoing global warming (Freudenburg & Muselli 2010).

Before running the models, all environmental variables were scaled to equal dimension and resolution (∼9km) and cropped to a rectangle with the extensions 70°W-30°W and 58°S-10°S, which encompass the skates’ current area of distribution. A Pearson’s correlation test was conducted to remove highly-correlated variables (|r| ≥ 0.8) from the analysis to avoid multicollinearity (Warren et al. 2014) and overfitting of models (Parolo et al. 2008). The ENMs for the current climatic scenario included the same variables selected after the Pearson correlation test for future modelling and followed the same procedures of scaling and cropping. The ENMeval package in R was used to partition occurrence data and select MaxEnt parameters (e.g., feature classes and regularization multiplier) to output the parsimonious model (ΔAICc = 0) (Muscarella et al. 2014). Localities were partitioned into model training and testing points applying the ‘block’ method, suitable when spatial or temporal transferability is required (Roberts et al. 2017).

### Measuring differences

We conducted a PCA-environment analysis (PCA-env) (Broennimann et al. 2012) to compare the niches per species in present and future climatic scenarios. Based on the classic Principal Component Analysis (PCA; Pearson 1901), PCA-env is an ordination approach to reduce into an uncorrelated linear combination of principal components (PCs) the climatic layers used in the ENMs. PCA-env was calibrated considering both climatic scenarios modelled so the position of occurrences along the PCs distinguishes between present and future environmental spaces (Broennimann et al. 2012). Resolution of the environmental grid was set to 100 pixels, and other parameters were kept to default.

We measured the niche overlap between present and future models per species by calculating the Schoener’s D index (Warren et al. 2008; Broennimann et al. 2012). Niche expansion, stability, and unfilling were also measured between intra-specific models of the two climatic scenarios (Guisan et al. 2014). Niche expansion is the portion of niche filled in a future climatic scenario, but not occupied in the current scenario; niche stability reflects the proportion of climatic conditions available in both temporal scenarios; and niche unfilling refers to conditions of current climatic scenario that are not filled in the projected future climatic scenario (Guisan et al. 2014).

Finally, niche similarity and equivalency were measured to test if present and future niches will be more similar and equivalent than expected at random for each species (Broennimann et al. 2012). In the former test, species present similar niches between the two scenarios if the measured similarity is significantly higher than randomly expected (*p* < 0.05), while in the latter, niches are considered equivalent if the measured value of equivalency falls within random expectations. In other words, for the niche equivalency test, significant results indicate that the niches are ecologically different. Both histograms of random similarity and equivalency scores are based on 100 repetitions of each test (Broennimann et al. 2012). All niche metrics were calculated using ade4 package version 1.7.13 and ecospat package version 3.0 in R (Chessel et al. 2004; Dray and Dufour 2007; Dray et al. 2007; Di Cola et al. 2017; Bougeard and Dray 2018).

## RESULTS

Six uncorrelated environmental variables were selected for the ecological niche models (ENMs) of each Riorajini species in current and future climatic scenarios: temperature mean (°C), salinity mean and range (psu), current velocity mean, minimum, and maximum (m^-1^). The most parsimonious models included different feature classes and values of regularization multiplier compared to default MaxEnt models (Table I).

**Table I.** Summary of the best combination (ΔAICc = 0) of parameters established by ENMeval package (Muscarella et al. 2014) per skate species and climatic scenario: P – present; F – future. n: number of occurrence points in the dataset; FC: Feature Classes allowed in the model (L – linear; Q – quadratic; H – hinge); RM: Regularization Multiplier; AUC: Average training Area Under ROC curve per model; sd: standard deviation of AUC.

| Species | Code | n |  | FC | RM | AUC | sd |
| --- | --- | --- | --- | --- | --- | --- | --- |
| *Atlantoraja castelnaui* | *Acas* | 31 | P | LQ | 0.5 | 0.986 | 0.003 |
|  |  |  | F | LQ | 1.5 | 0.982 | 0.003 |
| *Atlantoraja cyclophora* | *Acyc* | 60 | P | LQ | 0.5 | 0.991 | 0.001 |
|  |  |  | F | H | 2.5 | 0.991 | 0.001 |
| *Atlantoraja platana* | *Apla* | 30 | P | H | 2.5 | 0.977 | 0.004 |
|  |  |  | F | H | 4.0 | 0.977 | 0.005 |
| *Rioraja agassizii* | *Raga* | 36 | P | LQH | 2.5 | 0.978 | 0.003 |
|  |  |  | F | H | 2.0 | 0.981 | 0.003 |

ENMs show an increase in habitat suitability for the occurrence of *Atlantoraja castelnaui* (Figure 1) and *Rioraja agassizii* (Figure 2) along the latitudinal gradient they occupy, and in La Plata river mouth. For *A. cyclophora*, such an increase occurs more expressively at the Brazilian coast (Figure 3). There is a slight loss in environmental adequacy for the occurrence of *A. platana* near the coastline of Rio de Janeiro (23°S) but an overall increase in habitat suitability in deeper areas, still constrained to the continental shelf (Figure 4). This species showed an abiotic niche of higher stability in the group. Occurrence records plotted into the future modelled climatic scenario fall into areas of up to 2°C warmer than the present scenario, as predicted to be the highest increase in temperature for the RCP 8.5 climatic scenario.

**Figure 1.**
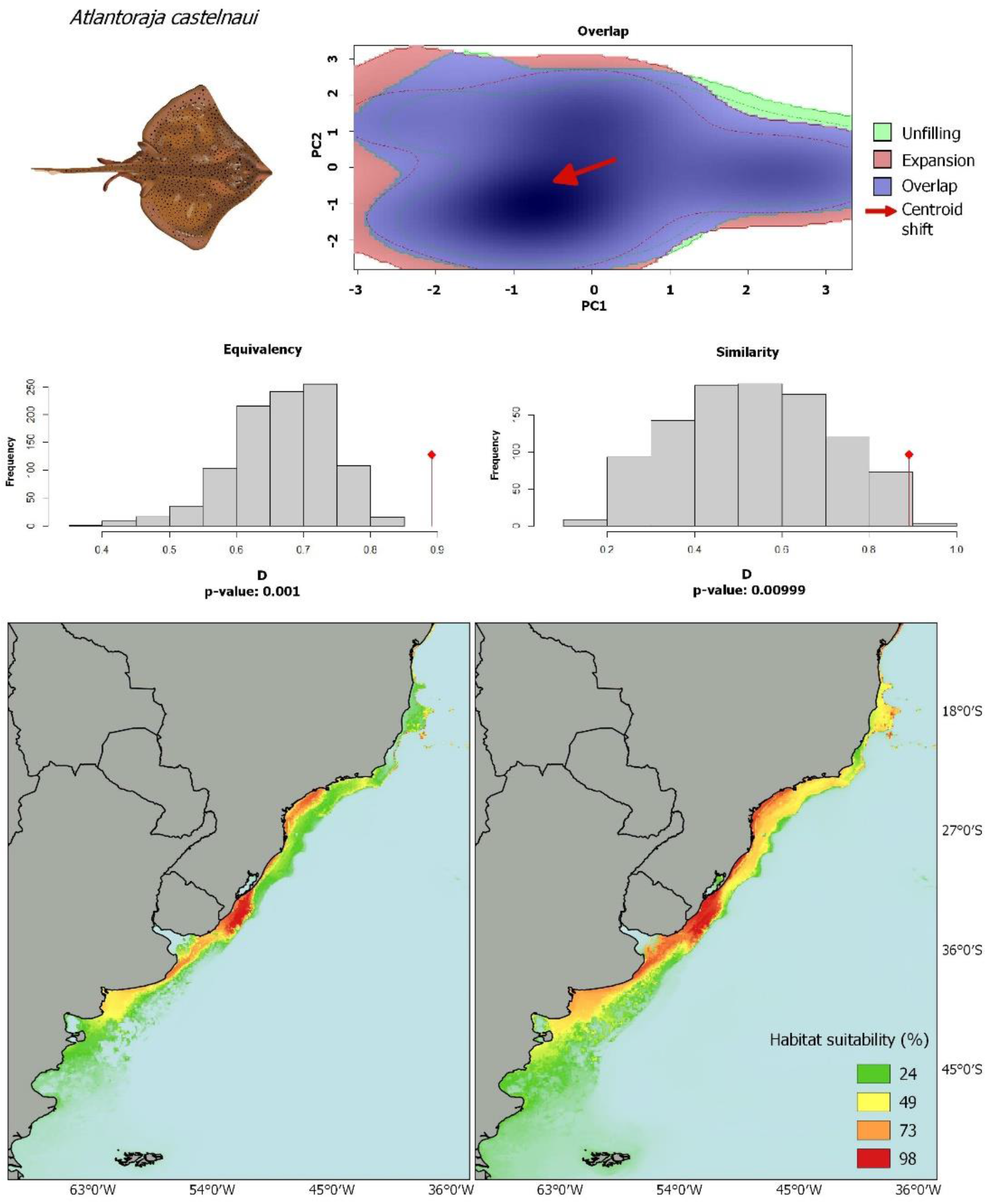
Niche dynamics and ecological niche models of present (left) and future (right) climatic scenarios showing percentage of environmental suitability for the occurrence of *Atlantoraja castelnaui*. The test of niche similarity indicates that the two climatic scenarios are more similar than randomly expected, although they are not equivalent (*p* < 0.05 in both similarity and equivalency tests).

**Figure 2.**
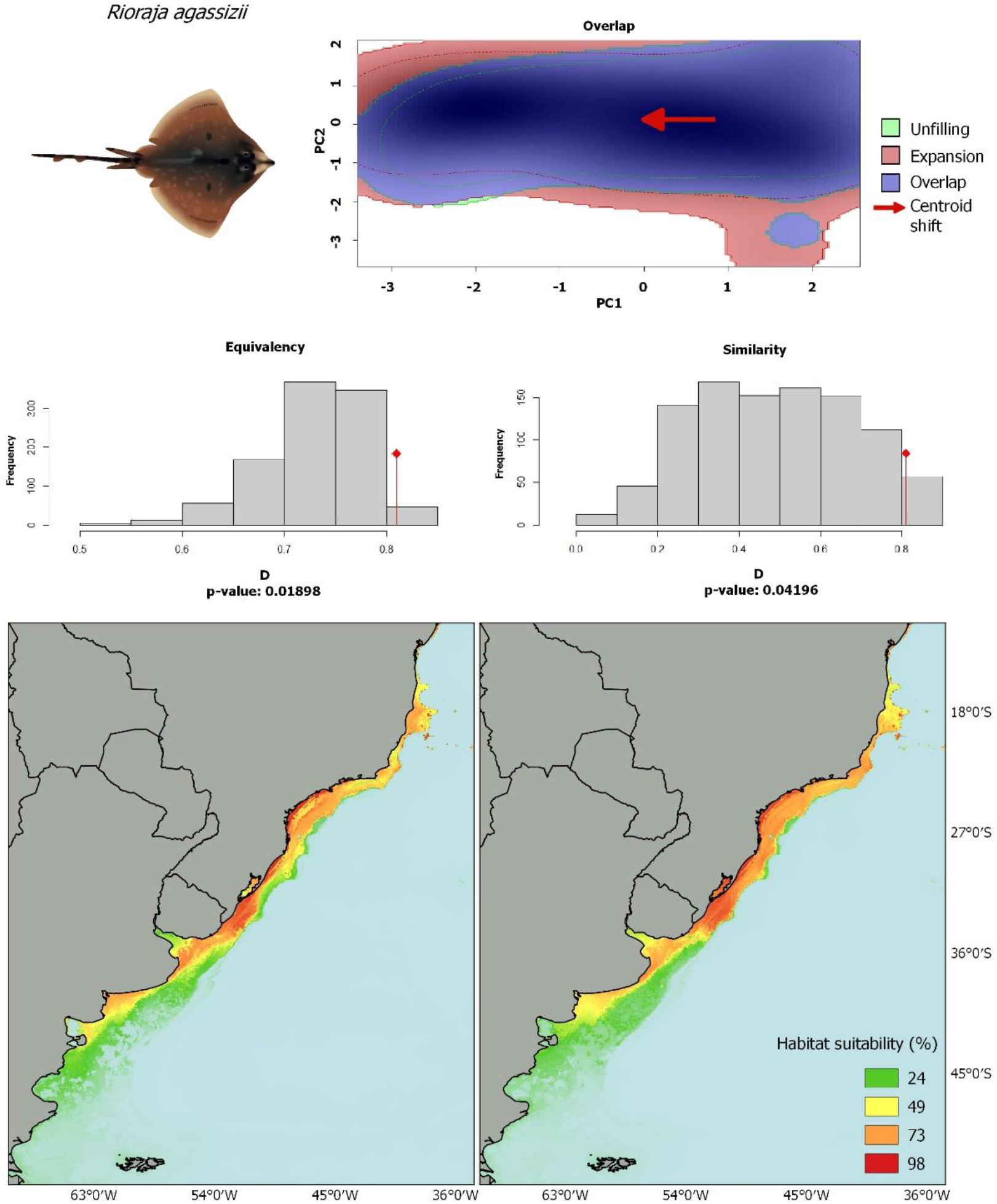
Niche dynamics and ecological niche models of present (left) and future (right) climatic scenarios showing percentage of environmental suitability for the occurrence of *Rioraja agassizii*. The test of niche similarity indicates that the two climatic scenarios are more similar than randomly expected, although they are not equivalent (*p* < 0.05 in both similarity and equivalency tests).

**Figure 3.**
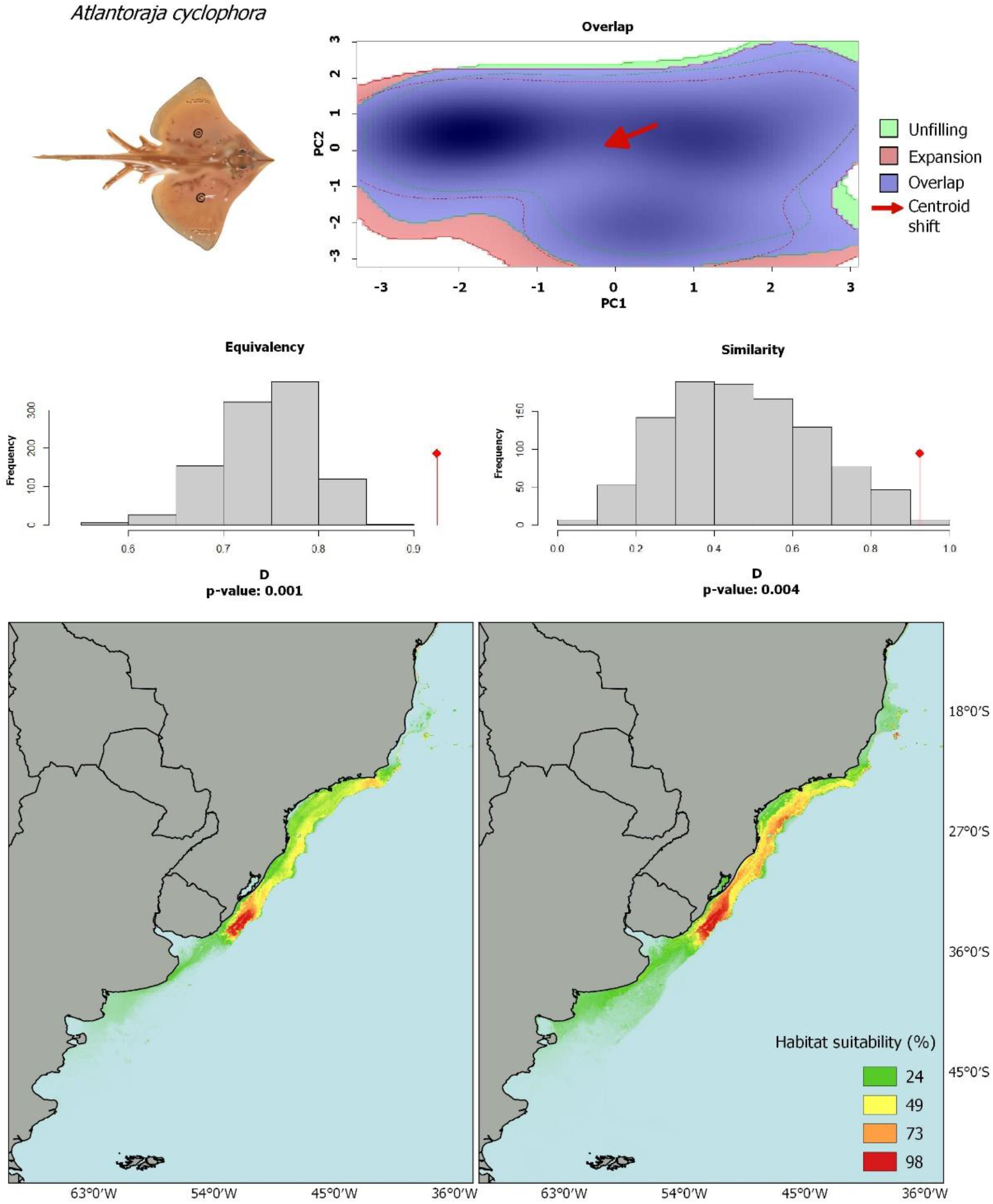
Niche dynamics and ecological niche models of present (left) and future (right) climatic scenarios showing percentage of environmental suitability for the occurrence of *Atlantoraja cyclophora*. The test of niche similarity indicates that the two climatic scenarios are more similar than randomly expected, although they are not equivalent (*p* < 0.05 in both similarity and equivalency tests).

**Figure 4.**
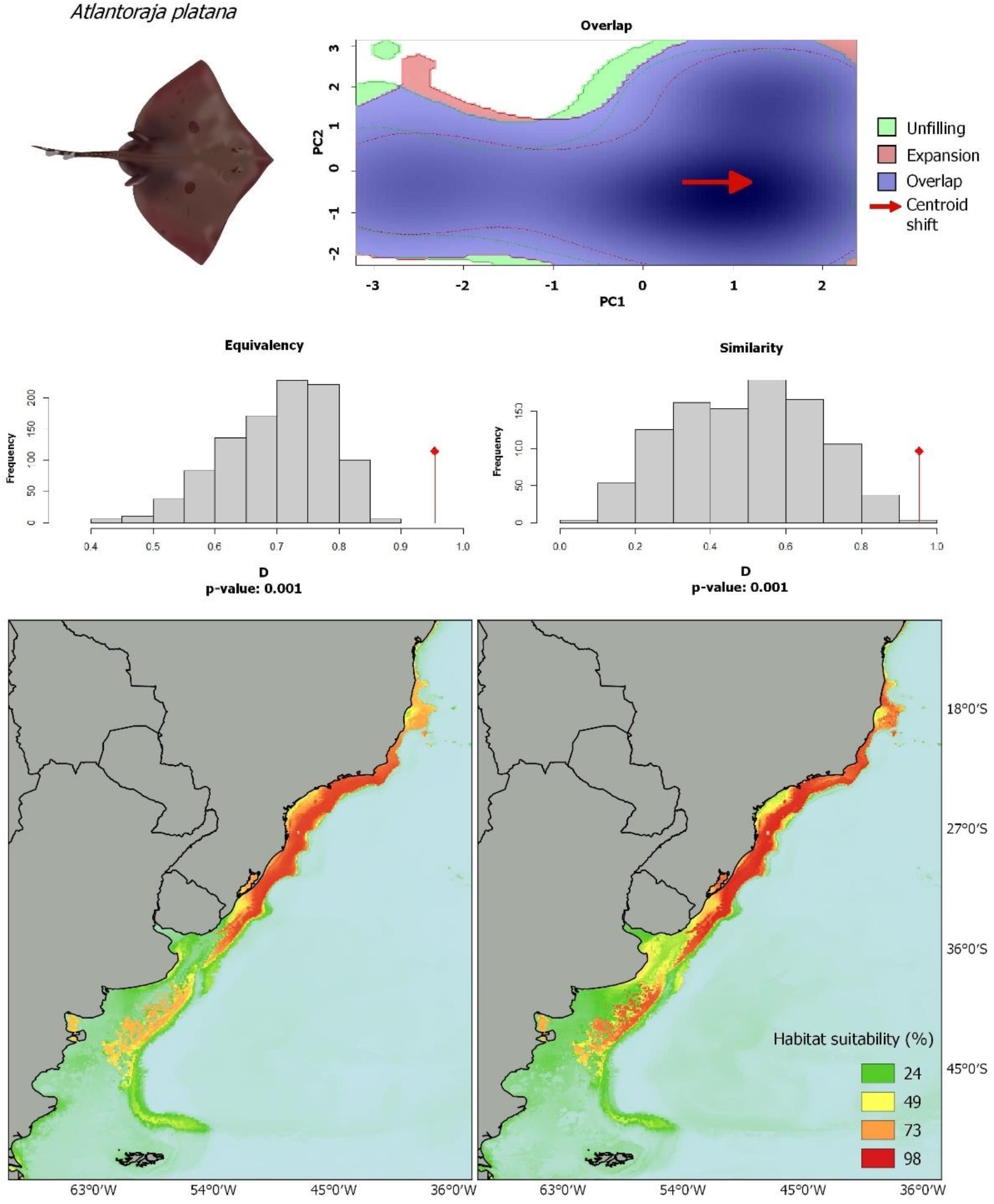
Niche dynamics and ecological niche models of present (left) and future (right) climatic scenarios showing percentage of environmental suitability for the occurrence of *Atlantoraja platana*. The test of niche similarity indicates that the two climatic scenarios are more similar than randomly expected, although they are not equivalent (*p* < 0.05 in both similarity and equivalency tests).

The importance of each of the six abiotic predictors included in the models varied in each climatic scenario modelled per species (Table II). Within species, niche overlap and stability between climatic scenarios were overall high (> 80%), and values of niche expansion and unfilling were low (< 22%), suggesting that the abiotic conditions currently required for the existence of these species in that area will be available in a future of warmer climatic conditions (Table III). Results of the niche similarity and equivalency tests indicate niches are similar albeit not equivalent between climatic scenarios (*p* < 0.05 in both tests for all species) (Figures 1–4). This means that in order to persist where they occur nowadays, under the worst-case scenario of climate change at temperate Southwestern Atlantic, Riorajini species should fill a niche that is similar but not equivalent to their current abiotic niche.

**Table II.**
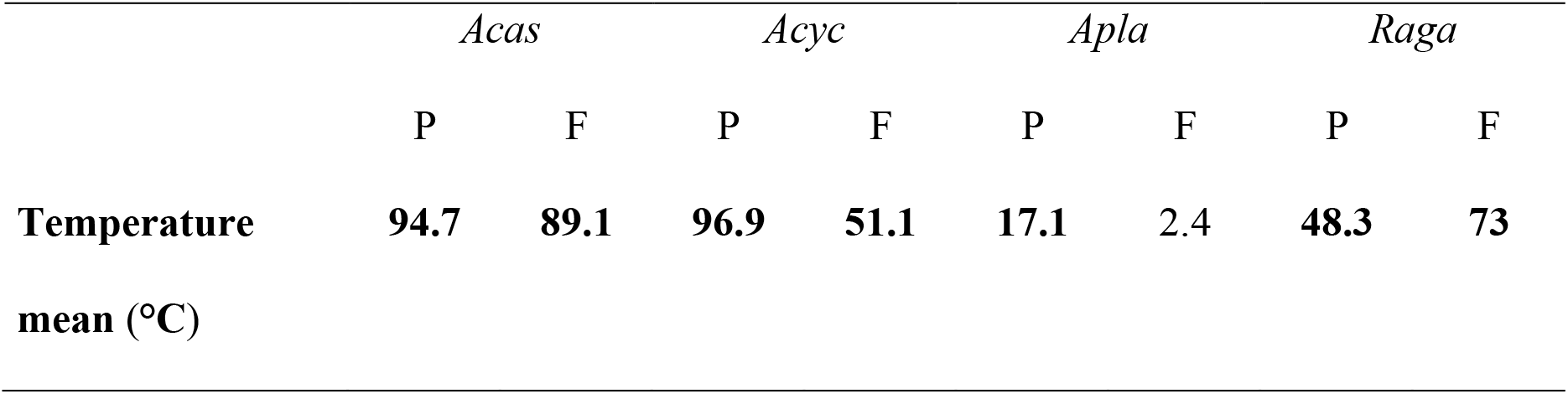
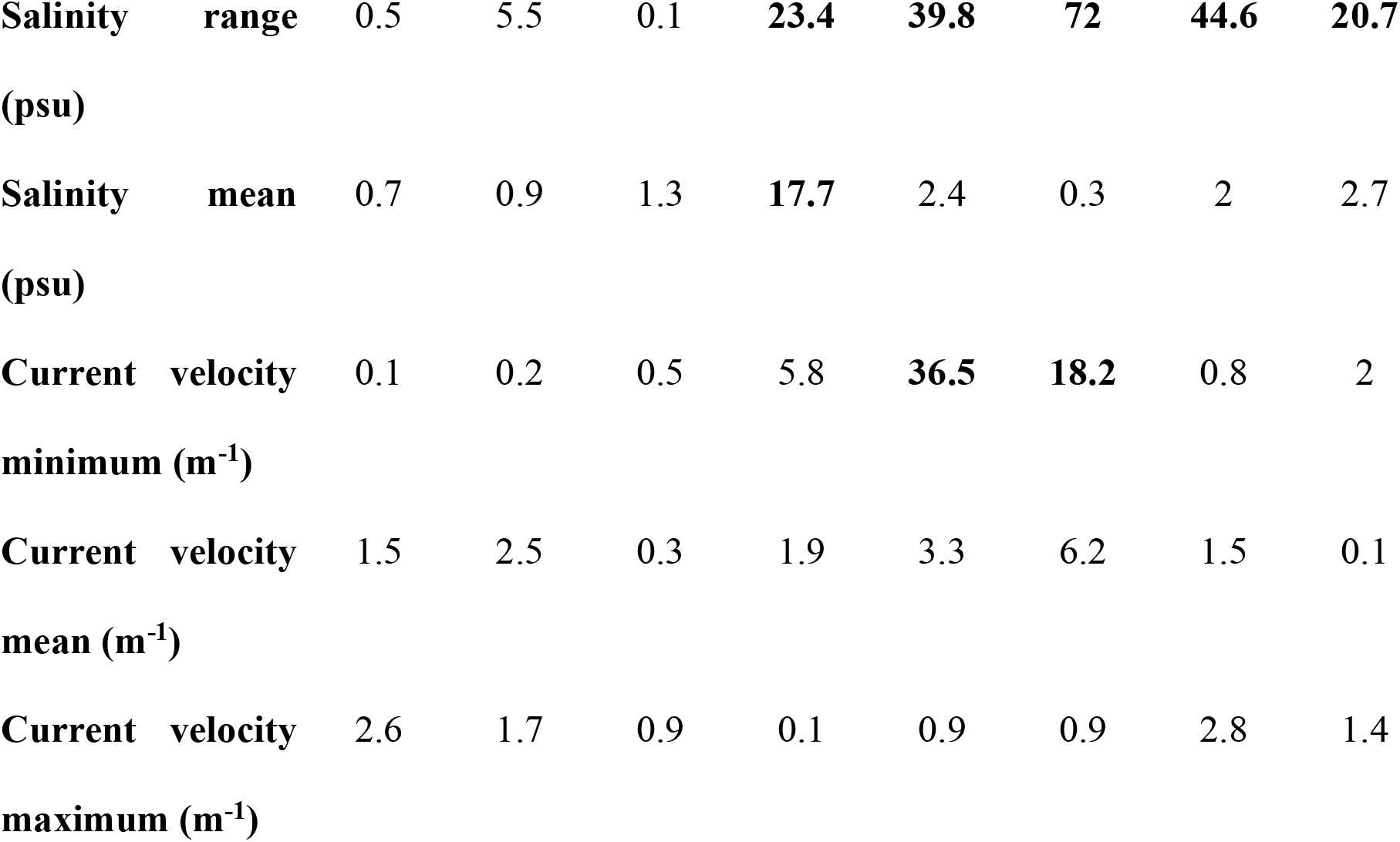
Permutation importance (%) per variable, per species for present (P) and future (F) climatic scenarios. *Acas* – *Atlantoraja castelnaui; Acyc* – *A. cyclophora*; *Apla – A. platana; Raga – Rioraja agassizii*. Bold highlights the variables of higher contribution (Σ > 80%) to models.

**Table III.** Niche overlap, expansion, stability, and unfilling measured between present and future climatic scenarios modelled for each Riorajini species. All values range from 0 (none) to 1 (maximum) and were rounded to two decimal places. *Acas* – *Atlantoraja castelnaui; Acyc* – *A. cyclophora*; *Apla – A. platana; Raga – Rioraja agassizii*.

|  | *Acas* | *Acyc* | *Apla* | *Raga* |
| --- | --- | --- | --- | --- |
| Niche overlap | 0.89 | 0.92 | 0.95 | 0.80 |
| Niche expansion | 0.19 | 0.10 | 0.05 | 0.10 |
| Niche stability | 0.80 | 0.90 | 0.94 | 0.90 |
| Niche unfilling | 0.21 | 0.07 | 0 | 0.10 |

## DISCUSSION

Temperate coastal zones sustain numerous social and economic activities of paramount importance to human communities like fisheries and ecotourism. Climate change is currently a major threat to the maintenance of these ecosystems’ goods and services along with its biodiversity (Pandolfi et al. 2003; Roessig et al. 2004; Pörtner & Peck 2010), making it crucial to identify areas of higher vulnerability and those of climatic stability that may pose as refugia to such impacts, in order to inform planning and implementing conservation efforts (Groves et al. 2012; Pacifici et al. 2015). In the Atlantic Southwest, the abiotic niche of endemic Riorajini skates tend to slightly expand in latitude and more markedly expand in longitude, towards deeper areas, as a consequence of climate change. This indicates that even the deeper layers of the water column where this group occurs are susceptible to dramatic environmental changes due to global warming, and this will potentially disturb interactions with other organisms. Overall, the southward range shift reduces areas for the occurrence of these species along the coast of Brazil, and increases in Uruguay and Argentina, thus demanding more international efforts for the conservation of this group.

A range shift towards higher depth in response to global warming, as seen in Riorajini, is the pattern for many other marine organisms, including invertebrates, teleosts, other elasmobranchs, and mammals, for example (Parmesan and Yohe 2003; Perry et al. 2005; Parmesan 2006; Molinos et al. 2015). Yet, here this shift occurs within the limits of the continental shelf, reinforcing the barrier that depth poses to the distribution of this group (Coelho et al. 2020). Among the species herein analyzed, *Rioraja agassizii* and the endangered species *Atlantoraja castelnaui* will face more changes because their abiotic niches presented the lowest values of overlap and highest values of niche unfilling between the climatic scenarios modelled, although they also presented the highest proportion of abiotic niche expansion (∼10 and ∼20%, respectively). *Atlantoraja cyclophora* and *A. platana*, on the other hand, presented the highest values of niche overlap between the two climatic scenarios (> 90% for both species), suggesting that the spots where these species currently occur will not face as severe changes.

Current abiotic conditions act like filters delimiting boundaries to the distribution of these taxa, but climate change effects could weaken such filters for aquatic invasive species (Rahel and Olden 2008). Nevertheless, the overall expansion of environmental adequacy does not necessarily translate into an organisms’ ability to occupy new climatically available areas, as other local forces might interact compromising dispersion (Vaz and Nabout 2016). VanDerWal et al. (2013) draw attention to the complexity of the combined climate change impacts and other factors influencing species distribution in a way so that simply looking at the expected poleward shift in biodiversity geographic distribution, detrimental to climate change underestimates the real effects of this phenomenon. Temperature mean was the variable of higher contribution to the models of present and future climatic conditions for all analyzed species except *A. platana*, which showed salinity range as the variable of higher contribution for both climatic scenarios (Table II). While higher temperatures might be tolerable for adults, it is likely to be harmful for young and eggs (Pörtner and Peck 2010). In addition to the physiological stresses imposed in young, cascading impacts from higher temperatures, such as ocean acidification and rise in sea level, directly threatens shallow-water habitats in coastal zones that are typical nursery areas for aquatic organisms (Roessig et al. 2004).

The role of temperature on the timing of hatching egg capsules of elasmobranchs is well documented in the literature (e.g., Clark 1922) and recent lab experiments have illustrated changes in their biology and physiology with probable link to global warming. For example, laboratory experiments simulating future concentrations of atmospheric carbon dioxide indicated behavioral alterations in sharks detrimental to water acidification (Green and Jutfelt 2014), as well as a decrease in metabolic and hunting efficiency (Pistevos et al. 2015). In skates, embryos of *Raja microocellata* Montagu, 1818 showed that increasing temperature in 2°C leads egg-capsules to hatch faster and produce young of 3.5% smaller body size (Hume 2019), and in embryos of *Leucoraja erinacea* (Mitchill, 1825), there is evidence of decrease in metabolic efficiency caused by both thermal stress and ocean acidification (Di Santo 2015). Such metabolic impacts in early developmental stages can reduce an organisms’ fitness and later compromise development and reproduction.

The lack of future projections for other environmental features in a context of climate change, as concentration of nitrate, a variable of high predictive power to the distribution of Riorajini (Coelho et al. 2020), was a limitation in our study. We somewhat addressed this caveat by comparing climatic scenarios modelled with the same environmental features. A broader assessment of climate change impacts in the biodiversity of a region should include various taxa of different life histories, and abiotic niches (e.g., species of algae, small invertebrates, top predators) to identify which groups are more vulnerable and visualize potential disruptions of ecological interactions from an ecosystem’ perspective. Future studies should take advantage of the increasing amount of biodiversity data available online (e.g., GBIF) and the numerous modelling approaches (e.g., Phillips et al. 2006; Broennimann et al. 2012; Guisan et al. 2014) to assess aspects of species’ biology and ecology in a relatively easy-to-follow, non-invasive frameworks, and inform putative conservation actions for threatened species in face of climate change. For the Riorajini group in particular, future modelling research should also include fisheries data to identify which populations are more exposed to these local pressures. This information is a crucial complement to the present study in order to inform plans of conservation efforts for these threatened species that considers local as well as large-scale impacts.

## CONCLUSION

Our study shows mainly a longitudinal increase in environmental suitability for the occurrence of four threatened Neotropical skates’ species in a projected scenario of global warming. This suggests that benthic layers of temperate coastal zones in the Atlantic Southwest are vulnerable to climate change even within a few decades used in this forecast. The shift in niche centroid between present and future climatic scenarios will push Riorajini species towards their limits of ecological tolerances and geographic space. Consequences of such shift can be detrimental both (i) to local biota in newly-occupied areas, as the introduction of a new predatory species can disturb the dynamics of this community, as well as (ii) to the species themselves, likely to face a reduction of nursery habitats in shallow waters due to an increasing heat. In summary, the niche expansion suggests that, under favorable biotic conditions, Riorajini species are likely to expand geographic range towards deeper zones although not surpassing the limits of the continental shelf. However, climate-related physiological stressors on skates’ eggs and young, and potential loss of nursery habitats may affect the reproductive success of these species and their ability to colonize new areas, which raises the question if an increase in environmental suitability will translate into actual occupancy.

## ACKNOWLEDGEMENTS

J. F. R. Coelho received financial support from CAPES (Coordenação de Aperfeiçoamento de Pessoal de Nível Superior). S. M. Q. Lima receives CNPq research productivity grant (313644/2018-7). We thank Maria Cristina Oddone, Juan Pablo Zurano, and Françoise Dantas de Lima for commenting on previous versions of the manuscript.

